# Temperature preference biases parental genome retention during hybrid evolution

**DOI:** 10.1101/429803

**Authors:** Caiti Smukowski Heil, Christopher R. L. Large, Kira Patterson, Maitreya J. Dunham

**Author notes:** Current address: National Heart, Lung, and Blood Institute, Bethesda, MD.

## Abstract

Interspecific hybridization can introduce genetic variation that aids in adaptation to new or changing environments. Here we investigate how the environment, and more specifically temperature, interacts with hybrid genomes to alter parental genome representation over time. We evolved *Saccharomyces cerevisiae* x *Saccharomyces uvarum* hybrids in nutrient-limited continuous culture at 15°C for 200 generations. In comparison to previous evolution experiments at 30°C, we identified a number of temperature specific responses, including the loss of the *S. cerevisiae* allele in favor of the cryotolerant *S. uvarum* allele for several portions of the hybrid genome. In particular, we discovered a genotype by environment interaction in the form of a reciprocal loss of heterozygosity event on chromosome XIII. Which species haplotype is lost or maintained is dependent on the parental species temperature preference and the temperature at which the hybrid was evolved. We show that a large contribution to this directionality is due to temperature sensitivity at a single locus, the high affinity phosphate transporter *PHO84*. This work helps shape our understanding of what forces impact genome evolution after hybridization, and how environmental conditions may favor or disfavor hybrids over time.

## Introduction

Comparative genomics of thousands of plants, animals, and fungi has revealed that portions of genomes from many species are derived from interspecific hybridization, indicating that hybridization occurs frequently in nature. However, the influence of processes such as selection, drift, and/or the presence or absence of backcrossing to a parental population on hybrid genome composition in incipient hybrids remains largely unknown. In some cases, hybrids will persist with both parental genomes in fairly equal proportions as new hybrid species or lineages, while in other instances, hybrid genomes will become biased towards one parent sub-genome over time [1–9]. Untangling the genetic and environmental factors that lead to these outcomes is a burgeoning field.

Some hybrid genotypes will be unfit due to genetic hybrid incompatibilities or cytotype disadvantage; decades of work across many systems have illustrated examples of hybrid sterility and inviability [10]. Recent work has demonstrated that in hybrid genomes with a bias in parental composition like humans, in which most of the genome is comprised of modern human haplotypes with small fragments derived from archaic human, regions from the minor parent (e.g., Neanderthal or Denisovan) are decreased near functional elements and hybrid incompatibilities [11–13]. Conversely, there are examples of “adaptive introgression,” in which alleles from the minor parent confer a benefit, like wing patterning in butterflies, high altitude tolerance in the Tibetan human population, and winter color morphs in the snowshoe hare [14–21]. The environment undoubtedly plays a significant role in hybrid fitness, and genotype by environment interactions will shape hybrid fitness in a similar manner as they shape non-hybrid fitness. For example, there is general acceptance that the *Saccharomyces* species complex is largely void of genic incompatibilities (with exceptions [22]), however most experiments looking for incompatibilities have used laboratory conditions. Hou *et al*. utilized different carbon sources, chemicals, and temperatures to show that over one-fourth of intraspecific crosses show condition-specific loss of offspring viability [23]. This is echoed by many examples of condition specific hybrid incompatibility in plants [24–29]. Similarly, there are numerous examples of environment dependent high fitness hybrid genotypes [30] [31–39], exemplified by classic research showing Darwin’s finch hybrids with different beak shapes gained a fitness benefit during and after an El Nino event [15].

The budding yeasts in the genus *Saccharomyces* have emerged as a particularly adept system to study genome evolution following hybridization. Recent evidence supports the hypothesis that the long-recognized whole genome duplication that occurred in the common ancestor that gave rise to *Saccharomyces* resulted from hybridization [40], and led to speculation that ancient hybridization could also explain other whole genome duplications in plants and animals [41]. Introgression and hybridization have also been detected across the *Saccharomyces* clade [42–48]; most famously, the lager brewing lineage *S. pastorianus* is a hybrid between *S. cerevisiae* and *S. eubayanus* [49–54]. A bias towards one parent sub-genome was identified in the ancient hybridization event and in *S. pastorianus*, and selection is inferred to be important in this process [1,40].

To empirically understand the genomic changes that occur as a hybrid adapts to a new environment, we previously created *de novo* interspecific hybrids between two yeast species, *S. cerevisiae* and *S. uvarum*, which are approximately 20 million years divergent and differ in a range of phenotypes, notably in preferred growth temperature. *S. uvarum* has been isolated from oak trees and associated soil in Patagonia and similar habitats across the world, and is specifically known for fermentation of cider and wines at cold temperatures [55–58]. Many *S.uvarum* strains show evidence of introgression from several other yeast species, and *S. cerevisiae* x *S. uvarum* hybrids have been isolated from fermentation environments [55,59].

We previously evolved *S. cerevisiae* x *S. uvarum* hybrids in the laboratory in several nutrient-limited environments at the preferred growth temperature of *S. cerevisiae* [60]. We frequently observed a phenomenon known as loss of heterozygosity (LOH) in these evolved hybrids, in which an allele from one species is lost while the other species’ allele is maintained. The outcome of such events is the homogenization of the hybrid genome at certain loci, and represents a way in which a hybrid genome may become biased toward one parent’s subgenome. This type of mutation can occur due to gene conversion or break induced repair, and as previously noted, has also been observed in organisms including *S. pastorianus*, pathogenic hybrid yeast, and hybrid plants, but its role in adaptation has been unclear [47,61,62]. We used genetic manipulation and competitive fitness assays to show that a particular set of LOH events was the result of selection on the loss of the *S. uvarum* allele and amplification of the *S. cerevisiae* allele at the high affinity phosphate transporter *PHO84* in phosphate limited conditions. By empirically demonstrating that LOH can be the product of selection, we illuminated how an underappreciated mutation class can underlie adaptive hybrid phenotypes.

This prior study illuminated how the environment (differences in nutrient availability) can bias a hybrid genome towards one parent sub-genome. Due to many examples of genotype by temperature interaction in hybrids across many taxa, and in particular difference in species temperature preference in our hybrids, we speculated that temperature is an important environmental modifier which may influence parental sub-genome representation in hybrids. Temperature can perturb fundamentally all physiological, developmental, and ecological processes, and as such, temperature is an essential factor in determining species distribution and biodiversity at temporal and spatial scales [63–65]. We hypothesized that in *S. cerevisiae* x *S. uvarum* hybrids, *S. cerevisiae* alleles may be favored at warmer temperatures, whereas *S. uvarum* alleles may be preferred at colder temperatures, giving the hybrid an expanded capacity to adapt. To test how temperature influences hybrid genome composition over time, we evolved the same interspecific hybrid yeast in the laboratory at 15°C for 200 generations. In comparing laboratory evolution at 15°C and 30°C, we present evidence that temperature can indeed bias hybrid genome composition towards one parental sub-genome, and we focus on a reciprocal LOH event at the *PHO84* locus. We show that which species’ allele is lost or maintained at this locus is dependent on the parental species’ temperature preference and the temperature at which the hybrid was evolved, thus revealing a genotype by environment interaction. Our results are one of the first clear examples with a molecular genetic explanation of how hybrids have expanded adaptive potential by maintaining two genomes, but also how adapting to one condition may abrogate evolutionary possibilities in heterogeneous environments.

## Results

### Laboratory evolution of hybrids and their parents at cold temperatures

To test whether temperature can influence the direction of resolution of hybrid genomes, we evolved 14 independent populations of a *S. cerevisiae* x *S. uvarum* hybrid in nutrient-limited media at 15° C for 200 generations (phosphate-limited: 6 populations; glucose-limited: 4 populations; sulfate-limited: 4 populations). Diploid *S. cerevisiae* and *S. uvarum* populations were evolved in parallel (4 populations of *S. cerevisiae* and 2 populations of *S. uvarum* in each of the three nutrient limited conditions). Populations were sampled from the final timepoint and submitted for whole genome sequencing and analysis.

### Loss of S. cerevisiae alleles in cold evolved hybrids

We detected large scale copy number variants in our cold evolved populations, including whole and partial chromosome aneuploidy and loss of heterozygosity (**Table 1**; **Tables S1-S2**; **Figures S1-S7**). Previously, in hybrids evolved at 30°C, we observed more LOH events in which the *S. uvarum* allele was lost (5/9 LOH events), and found a significant preference for *S. cerevisiae* partial and whole chromosome amplification [60]. In contrast, in hybrids evolved at 15°C, we observe 6/6 LOH events in which the *S. cerevisiae* allele is lost and the *S. uvarum* allele is maintained, suggestive of a *S. uvarum* cold temperature benefit. While our sample sizes are modest, together these results indicate that temperature can determine hybrid genomic composition in the generations following a hybridization event.

**Table 1:** Mutations in cold-evolved hybrid populations.

| Population | Location | Gene(s) | Mutation |
| --- | --- | --- | --- |
| **P1-15°** | *S. cerevisiae* chrXVI: 772650 | *CLB2* | coding-nonsynonymous: D333A |
|  | *S. uvarum* chrVII: 818275 | *UBR1* | coding-nonsynonymous: F1634C |
|  | *S. uvarum* chrXI: 5832 | *JEN1* | 5'-upstream |
|  | *S. cerevisiae* chrXIII: 1-81102<br>*S. uvarum* chrXIII: 1-82283 | 41 genes including *PHO84* | LOH: loss of *S. cerevisiae* allele and amplification of *S. uvarum* allele |
|  | *S. cerevisiae* chrXIII: 81102-168343 | 47 genes | CNV: Segmental amplification of *S. cerevisiae* |
| **P2-15°** | *S. uvarum* chrIV: 903941 | *YBR259W* | coding-nonsynonymous: S624Y |
| **P2F-15°** | *S. uvarum* chrV: 179951 | *YAT2* | coding-synonymous |
| **P3-15°** | *S. uvarum* chrXI: 101238 | *LST4* | coding-nonsynonymous: S533A |
|  | *S. cerevisiae* chrXIII: 1-79085 | 40 genes including *PHO84* | LOH: loss of *S. cerevisiae* allele and amplification of *S. uvarum* allele |
|  | *S. uvarum* chrXIII:1-80181 |  |  |
|  | *S. cerevisiae* chrXIII:79085-168343 | 48 genes | Segmental amplification of *S. cerevisiae* |
| **G7-15°** | *S. cerevisiae* chrXI:80553 | *ACP1* | 5' upstream |
|  | *S. cerevisiae* chrIV:651345-871844 | 118 genes | Segmental amplification *S. cerevisiae* allele |
|  | *S. uvarum* chrII:554234-1289935 | 159 genes | Segmental amplification *S. uvarum* allele |
|  | *S. cerevisiae* chrXV:976083-1071297 | 46 genes | LOH: loss of *S. cerevisiae* allele |
| **G8-15°** | *S. cerevisiae* chrIV:955826 | *VHS1* | 5'-upstream |
|  | *S. cerevisiae* chrV:321068 | *AIM9* | coding-nonsynonymous: A369V |
|  | *S. cerevisiae* chrXI:80553 | *CCP1* | 5'-upstream |
|  | *S. cerevisiae* chrXV:977571 | *TYE7* | coding-nonsynonymous: E167K |
|  | *S. uvarum* chrXV:685069 | *IKI1* | coding-nonsynonymous: A2E |
|  | *S. cerevisiae* chrIII:169495-316620 | 82 genes | LOH: loss of *S. cerevisiae* allele |
|  | *S. cerevisiae* chrXII:732111-1078177 | 181 genes | Segmental amplification of *S. cerevisiae* allele |
|  | *S. cerevisiae* chrXV: 340969-464306,464306-594878 | 71 genes, 72 genes | LOH: loss of *S. cerevisiae* allele |
| **G9-15°** | *S. cerevisiae* chrXII:812389 | *FKS1* | coding-nonsynonymous: S798Y |
|  | *S. cerevisiae* chrXII:818575-1078177 | 136 genes | Segmental amplification of *S. cerevisiae* allele |
|  | *S. cerevisiae* chrXV:976083-1071297 | 46 genes | LOH: loss of *S. cerevisiae* allele |
| **G10-15°** | *S. cerevisiae* chrXV:1034298 | *GPB1* | 5' upstream |
| **S7-15°** | *S. cerevisiae* chrXII:238463 | *YLR046C* | coding-nonsynonymous: M117I |
|  | *S. cerevisiae* chrXII:1047895 | *FMP27* | coding-nonsynonymous: P1300S |
|  | *S. cerevisiae*<br>chrII:786030-813184 | 12 genes<br>including *SUL1* | Segmental amplification of *S. cerevisiae* allele |
| **S8-15°** | *S. cerevisiae*<br>chrIX:212119 | *AIR1* | 5' upstream |
|  | *S. cerevisiae*<br>chrIX:212130 | *AIR1* | 5' upstream |
|  | *S. cerevisiae*<br>chrII:786025-813184 | 12 genes<br>including *SUL1* | Segmental amplification of *S. cerevisiae* allele |
| **S9-15°** | *S. cerevisiae*<br>chrII:786017-813184 | 12 genes<br>including *SUL1* | Segmental amplification of *S. cerevisiae* allele |
| **S10-15°** | *S. uvarum* chrXIII:<br>752118 | *GTO3* | 5' upstream |
|  | *S. uvarum*<br>chrXVI: 456125 | *SUR1* | 5' upstream |
|  | *S. cerevisiae*<br>chrII:770240-813184 | 21 genes<br>including *SUL1* | Segmental amplification of *S. cerevisiae* allele |

LOH: loss of heterozygosity; CNV: copy number variant. No mutations were detected in populations P4-15°, P5-15°, or P6-15°. Breakpoints of CNV and LOH are approximate.

Copy number changes, and specifically amplification of nutrient transporter genes, are well-recognized paths to adaptation in laboratory evolution in nutrient limited conditions [48,60,66–75]. Similar to previous studies, we find both chromosomal aneuploidy and LOH are nutrient limitation specific, with repeatable genomic changes occurring in replicate populations under the same nutrient condition, but few if any changes shared across nutrients. In glucose limitation, 3/4 hybrid populations experienced chromosome XV LOH, losing the *S. cerevisiae* allele for portions of the chromosome. Haploidization of one of these implicated regions on chromosome XV was previously observed in *S. cerevisiae* diploids evolved at 30°C in glucose limitation [60,73], but it was not observed in any previously evolved hybrids, and which genes may be responsible for fitness increases are unclear. In sulfate limitation, we recapitulate previous hybrid laboratory evolution results [60], observing the amplification of the *S. cerevisiae* high affinity sulfate transporter gene *SUL1* in low sulfate conditions (4/4 hybrids, **Figures S1, S2**). Amplification of *S. cerevisiae SUL1* therefore seems to confer a high relative fitness regardless of temperature (see section below, **“**Pleiotropic fitness costs resulting from loss of heterozygosity”). Though prior work showed highly repeatable amplification of *S. cerevisiae SUL1* at 30°C in *S. cerevisiae* haploids and diploids [60,66,73–75], and amplification of *S. uvarum SUL2* after approximately 500 generations at 25°C in *S. uvarum* diploids [75], we never observed amplification of *SUL1* or *SUL2* in *S. cerevisiae* or *S. uvarum* diploids at 15°C, albeit our experiments were terminated at 200 generations.

Finally and most notably, in low phosphate conditions, we discovered a LOH event in which the *S. cerevisiae* allele is lost and *S. uvarum* allele is amplified on chromosome XIII, which encompasses the high affinity phosphate transporter *PHO84* locus (2/6 hybrid populations; **Figure 1 A**). The directionality of this LOH event is the opposite outcome of our observations of hybrids evolved at 30° C, in which the *S. cerevisiae* allele was amplified and the *S. uvarum* allele was lost in this same region (3/6 populations, **Figure 1 A**). The direction of resolution of these LOH events thus appears to be modulated by temperature.

**Figure 1:**
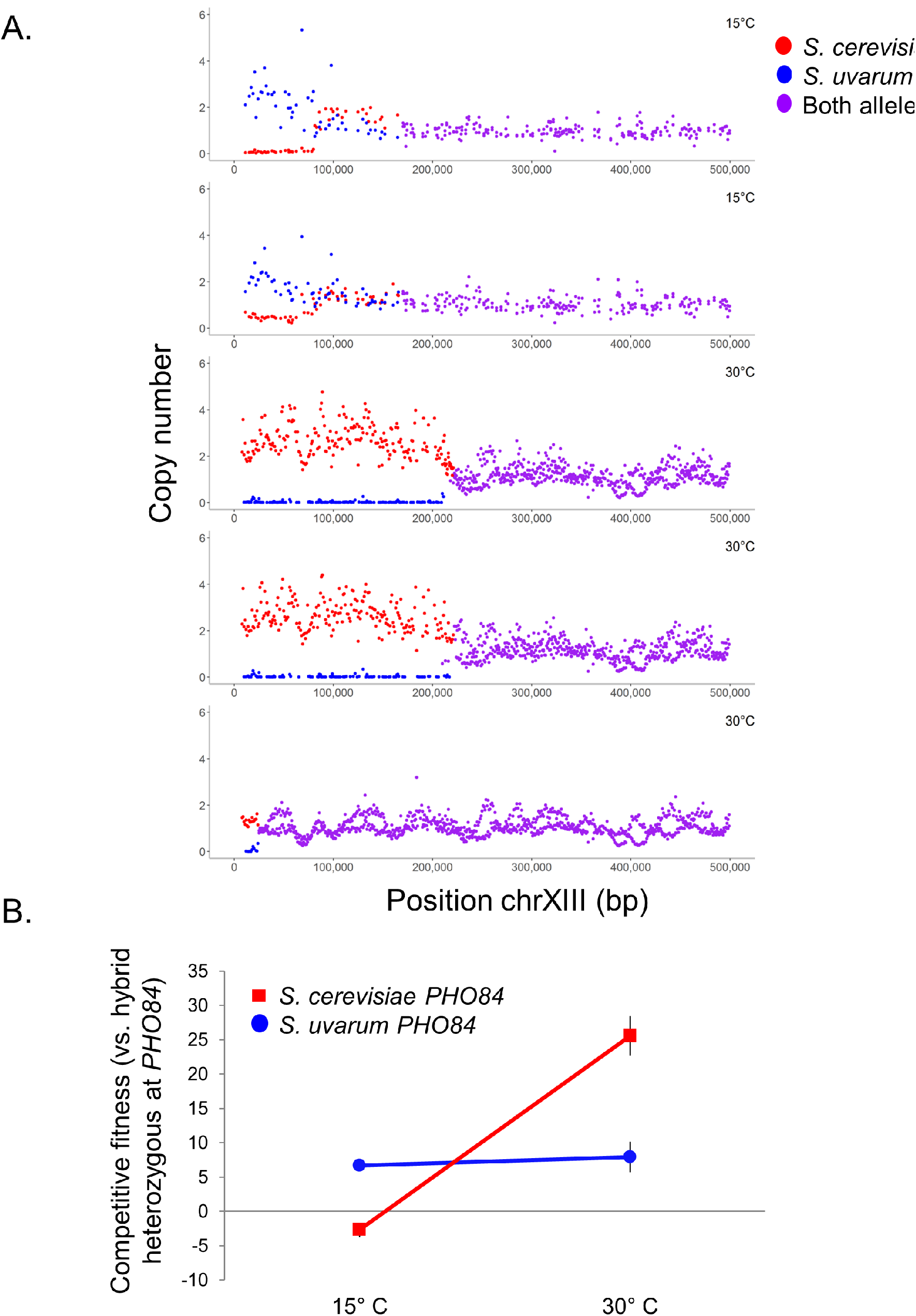
Loss of heterozygosity directionality results from selection on different species’ alleles at different temperatures. **A**. Evolved hybrids exhibit reciprocal loss of heterozygosity on chromosome XIII encompassing the high affinity phosphate transporter gene *PHO84* in phosphate limited conditions at different temperatures. 2/6 independent populations lost the *S. cerevisiae* allele when evolved at 15°C (top 2 panels), 3/6 independent populations lost the *S. uvarum* allele when evolved at 30°C (bottom 3 panels). Purple denotes a region where both alleles are present at a single copy, blue denotes a *S. uvarum* change in copy number, red denotes a *S. cerevisiae* change in copy number. Note, copy number was derived from sequencing read depth. Clone sequencing was utilized for experiments at 30°C and population sequencing was utilized for experiments at 15°C, so exact population frequency and copy number changes are unclear for experiments at 15°C. **B**. Allele swap experiments in which a hybrid with one allele of *PHO84* from each species is competed against a hybrid with both copies of *PHO84* either from *S. cerevisiae* (Sc*PHO84*/Sc*PHO84*; red) or *S. uvarum* (Su*PHO84*/Su*PHO84*; blue) reveal a fitness effect dependent on temperature.

### Environment-dependent loss of heterozygosity aids in temperature adaptation in hybrids

Based on previous results that demonstrated that LOH at the *PHO84* locus conferred a high fitness benefit at warm temperatures, we hypothesized that this apparent preference for the alternate species’ allele in different environments is explained by a genotype by environment interaction at the *PHO84* locus itself. To test this hypothesis, we repeated the competitive fitness assays of allele-swapped strains from Smukowski Heil *et al*. (2017) at 15°C. These strains are either homozygous *S. cerevisiae*, homozygous *S. uvarum*, or heterozygous for both species at the *PHO84* locus in an otherwise isogenic hybrid background. Indeed, we demonstrate a fitness tradeoff dependent on temperature, in which hybrids homozygous for *S. uvarum PHO84* show a fitness increase of 6.69% (+/−0.49) at 15°C relative to their hybrid ancestor, which carries a copy of each species’ *PHO84* allele. In contrast, hybrids homozygous for *S. cerevisiae PHO84* show a slight relative fitness decrease (-2.67% +/−1.00) at this temperature (**Figure 1 B**). There is a significant difference between fitness of hybrids homozygous for *S. cerevisiae PHO84* at different temperatures (p<0.001, Welch Two Sample t-test), suggesting the *S. cerevisiae* allele of *PHO84* is temperature sensitive.

### Pleiotropic fitness costs resulting from loss of heterozygosity

We clearly demonstrate a fitness trade-off dependent on temperature at the *PHO84* locus. To explore if other mutations in evolved hybrids demonstrate antagonistic pleiotropy at divergent temperatures, we conducted a series of competitive fitness assays at 15°C and 30°C. We isolated two clones from each hybrid population evolved at 15°C, and competed the clone against a common unevolved hybrid ancestor in the nutrient limitation it was evolved in at both 15°C and 30°C.

First, we sought to identify how the chromosome XIII LOH event influences fitness beyond the *PHO84* locus. We demonstrate that clones evolved in phosphate limitation with the chromosome XIII LOH event (homozygous *S. uvarum PHO84*; P1–15°C and P3–15°C) have higher competitive fitness at 15°C and decreased competitive fitness at 30°C, displaying antagonistic pleiotropy (**Figure 2 A**). In contrast, clones isolated from populations without chromosome XIII LOH (P2–15°C, P4–15°C, P5–15°C, P6–15°C) have variable fitness responses at both temperatures. To compare these results to the reciprocal LOH event seen in hybrids evolved at 30°C in which hybrids became homozygous for *S. cerevisiae PHO84*, we competed clones from populations initially evolved at 30°C at 15°C. Indeed, clones with the LOH event homozygous for *S. cerevisiae PHO84* (P3–30°C, P4–30°C, P5–30°C) have increased fitness at 30°C and decreased fitness at 15°C (**Figure 2 B**), consistent with the *PHO84* allele swap competitive fitness results. Of course, there are other mutations present in these clones, and some evidence that these fitness values may be influenced by the tract length of the LOH event, which ranges from approximately 79kb to 234kb. For example, P3–30°C has the shortest LOH tract at approximately 25kb in length and has a higher relative fitness at 15°C than either P4–30°C or P5–30°C, whose LOH tracts extend to 221kb and 234kb, respectively. The LOH tract length is approximately 80kb in both cold evolved populations (P1–15°C: 82,283; P3–15°C: 79,085), but is made more complex by the amplification of a portion of the *S. cerevisiae* sub-genome adjacent to the LOH event (P1–15°C: 81,105–168,345; P3–15°C: 79,074–168,345; **Figure 1 A**). Together, these results support a temperature sensitive fitness response at the *PHO84* locus, but also imply that there may be other genes modulating fitness in the chromosome XIII LOH events, something we hope to explore in future work.

**Figure 2:**
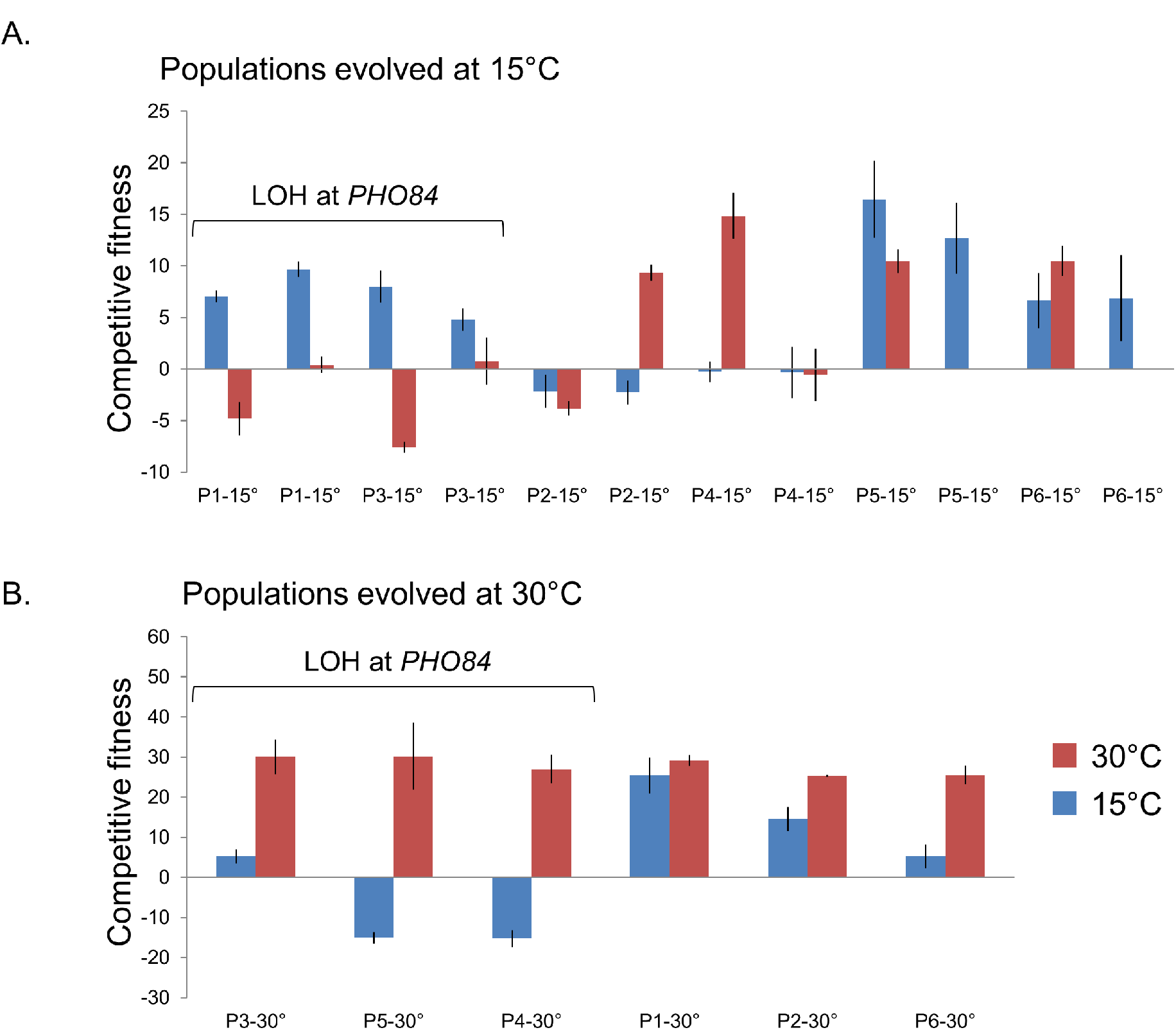
Fitness assays demonstrate that loss of heterozygosity results in antagonistic pleiotropy. **A**. One or two clones were isolated from each population evolved in phosphate limitation at 15°C and competed against a common competitor, the hybrid ancestor of the evolution experiments, at 15°C (blue) and 30°C (red) in phosphate limitation. Clones with chromosome XIII loss of heterozygosity exhibited higher fitness relative to their ancestor at 15°C and neutral or negative fitness at 30°C. Error bars denote standard error from technical and/or biological replicates. **B**. Clones evolved in phosphate limitation at 30°C were competed against a common competitor, the hybrid ancestor of the evolution experiments, at 15°C and 30°C in phosphate limitation. Data from 30°C fitness assays was obtained from [60]. Clones with chromosome XIII loss of heterozygosity exhibited higher fitness relative to their ancestor at 30°C and neutral or negative fitness at 15°C.

Results are variable from hybrid clones evolved in other media conditions at 15°C, with some clones having higher relative fitness at 15°C and lower fitness at 30°C, some clones showing the opposite trend, and some clones having similar fitness at both temperatures (**Table 2**). It thus appears that temperature specific antagonistic pleiotropy, in which a clone has high fitness at one temperature and low fitness at the other temperature, is relatively rare, with the LOH encompassing *PHO84* being the only clear example (but see P2-C2, G9-C2). The only other distinct pattern in the fitness data is that all hybrids evolved in sulfate limitation demonstrate fitness gains at both 15°C and 30°C. All populations have an increased fitness of 23.66–41.01% relative to their hybrid ancestor at 30°C, except for the clone from population S9–15°C. This result is in line with the observation of an amplification of *SUL1* at very low frequency and/or low copy number in this population compared to other sulfate limited evolved populations (**Figure S1**). These data suggest that an amplification of *S. cerevisiae SUL1* confers a fitness benefit at both cold and warm temperatures, but is most beneficial at warm temperatures.

**Table 2:** Competitive fitness of hybrids at two temperatures.

| Population Identifier | Original evolution conditions | Relative Fitness at 15°C (+/-S.E.) | Relative Fitness at 30°C (+/- S.E.) |
| --- | --- | --- | --- |
| G7-15° | Glucose limitation, 15°C | C1: -0.66<br>C2: 6.37 (+/-3.96) | C1: 1.86<br>C2: 6.63 (+/-1.39) |
| G8-15° | Glucose limitation, 15°C | C1: 2.65 (+/-0.85)<br>C2: 14.83 (+/-3.63) | C2: 2.07 (+/-3.91) |
| G9-15° | Glucose limitation, 15°C | C1: 6.40 (+/-0.28)<br>C2: 11.76 | C1: 10.10 (+/-2.83)<br>C2: -2.45 (+/-1.68) |
| G10-15° | Glucose limitation, 15°C | C1: 5.14 (+/-0.03) | C1: 13.06 (+/-4.60) |
| S7-15° | Sulfate limitation, 15°C | C1: 7.43 (+/-0.27)<br>C2: 28.52 (+/-1.42) | C1: 23.66<br>C2: 30.12 (+/-10.20) |
| S8-15° | Sulfate limitation, 15°C | C1: 18.55 (+/-3.98)<br>C2: 30.85 (+/-1.18) | C1: 39.38 (+/-1.25)<br>C2: 37.47 (+/-0.77) |
| S9-15° | Sulfate limitation, 15°C | C1: 7.40 (+/-4.34) | C1: 12.54 (+/-3.70) |
| S10-15° | Sulfate limitation, 15°C | C1: 12.26 (+/-1.26) | C1: 41.01 (+/-4.03) |
| P1-15° | Phosphate limitation, 15°C | C1: 7.05 (+/-0.59)<br>C2: 9.68 (+/-0.76) | C1: -4.81 (+/-1.62)<br>C2: 0.40 (+/-0.79) |
| P2-15° | Phosphate limitation, 15°C | C1: -2.16 (+/-1.60)<br>C2: -2.26(+/-1.16) | C1: -3.82 (+/-0.69)<br>C2: 9.34 (+/-0.76) |
| P3-15° | Phosphate limitation, 15°C | C1: 8.01 (+/-1.55)<br>C2: 4.80 (+/-1.07) | C1: -7.59 (+/-0.52)<br>C2: 0.76 (+/-2.30) |
| P4-15° | Phosphate limitation, 15°C | C1: -0.26 (+/-1.00)<br>C2: -0.33 (+/-2.51) | C1: 14.87 (+/-2.21)<br>C2: -0.58 (+/-2.55) |
| P5-15° | Phosphate limitation, 15°C | C1: 16.46 (+/-3.73)<br>C2: 12.69 (+/-3.44) | C1: 10.48 (+/-1.16) |
| P6-15° | Phosphate limitation, 15°C | C1: 6.66 (+/-2.66)<br>C2: 6.90 (+/-4.15) | C1: 10.49 (+/-1.45) |
| P1-30° | Phosphate limitation, 30°C | 21.01 | 29.18 (+/-1.37) |
| P2-30° | Phosphate limitation, 30°C | 14.55 (+/-2.94) | 25.34 (+/-0.24) |
| P3-30° | Phosphate limitation, 30°C | 5.25 (+/-1.77) | 30.03 (+/-4.31) |
| P5-30° | Phosphate limitation, 30°C | -8.14 (+/-0.81) | 30.24 (+/-8.32) |
| P4-30° | Phosphate limitation, 30°C | -10.98 (+/-2.35) | 27.02 (+/-3.62) |
| P6-30° | Phosphate limitation, 30°C | -2.06 (+/-0.66) | 25.52 (+/-3.32) |

One or two clones (denoted as C1, C2) were selected from each cold-evolved population and competed against a GFP-tagged ancestor in the nutrient limitation they in which they were evolved at both 15°C and 30°C. Six clones evolved in phosphate limitation at 30°C from Smukowski Heil et al. 2017 were also tested at 15°C (fitness data at 30°C in [60]).

### *Single nucleotide variants show no overlap with previous study, occurrence shows slight* S. cerevisiae *sub-genome bias in hybrids*

Through comparison of the single nucleotide variants and indels called in the hybrid populations evolved at 15°C and 30°C, we observed a slight, though not significant, increase in the number of mutations in the *S. cerevisiae* portion of the genome when evolved at temperatures preferred by *S. uvarum* (12/19 mutations are in the *S. cerevisiae* sub-genome at 15°C compared to 16/30 mutations in the *S. cerevisiae* sub-genome at 30°C, Fisher’s exact test p = 0.5636; **Table S3**). We speculate that the relationship between the number of detectable, high-frequency mutations and growth temperature could be reflective of the degree to which either species is shifted from its optimal environment, and a difference in the size of the adaptive space available. There was no overlap in genes with variants identified in datasets from 15°C and 30°C.

We suspect that the low growth temperature is a selective pressure for both the hybrids and the parental populations, and we did observe mutations in two genes (*BNA7* and *OTU1*) that were previously identified in a study of transcriptional differences of *S. cerevisiae* in long-term, glucose-limited, cold chemostat exposure [76]. We found no overlap with genes previously identified to be essential for growth in the cold [77,78], or differentially expressed during short-term cold exposure [79,80], though our screen is hardly saturated and growth conditions differ between these studies. Additionally, we observed some mutations in genes that are members of the cAMP-PKA pathway, which has been previously implicated in cold and nutrient-limitation adaptation [76,81].

Based on the mutations observed in populations evolved at 30°C, we previously hypothesized that an intergenomic conflict between the nuclear and mitochondrial genome of *S. cerevisiae* and *S. uvarum* is an important selection pressure during the evolution of these hybrids [60]. We find further circumstantial evidence for the possibility that mitochondrial conflicts are influential in hybrid evolution as 3/19 point mutations in the hybrids are related to mitochondrial function, whereas 1/20 are related in the parental species populations.

Finally, we did observe one recurrent mutational target. Eight independent *S. cerevisiae* diploid lineages had a substitution occur at 1 of 3 different amino acid positions in Tpk2, a cAMP-dependent protein kinase catalytic subunit. Previously, it has been reported that Tpk2 is a key regulator of the cell sticking phenotype known as flocculation through inactivation of Sfl1, a negative regulator of *FLO11*, and activation of *FLO8*, a positive regulator of *FLO11* [82,83]. This mutation was detected exclusively in flocculent populations. We and others have previously established that flocculation evolves quite frequently in the chemostat, likely as an adaptation to the device itself, but we have not previously observed a flocculation phenotype caused by these mutations in other evolved populations of *S. cerevisiae* [84]. While we have not definitively demonstrated causation, prior literature links *TPK2* to flocculation, and all evolved clones bearing a *TPK2* mutation flocculated within seconds of resuspension by vortexing (**Figure S8**). Most mutations were heterozygous, but within several lineages, we observed evidence of a LOH event that caused the *TPK2* mutation to become homozygous. Clones bearing a homozygous mutation in *TPK2* showed a faster flocculation phenotype than their heterozygous counterparts. We similarly observed one lineage with a mutation causing a pre-mature stop and subsequent LOH in *SFL1*, whose isolated clones displayed a robust flocculation phenotype. We suspect that our previous lack of detection is likely due to the well-established genetic differences in the *FLO8* gene between the strains used in this study and previous studies, which would alter whether a *FLO8* dependent flocculation phenotype is possible [85].

## Discussion

In summary, we evolved populations of interspecific hybrids at cold temperatures and show that temperature can influence parental representation in a hybrid genome. We find a variety of mutations whose annotated function is associated with temperature or nutrient limitation, including both previously described and novel genes. Most notably, we discover a temperature and species specific gene by environment interaction in hybrids, which empirically demonstrates that temperature influences hybrid genome evolution.

Growth temperature appears to be one of the most definitive phenotypic differences between species of the *Saccharomyces* clade, with *S. cerevisiae* being exceptionally thermotolerant, while many other species exhibit cold tolerance [86–88]. Significant work has focused on determining the genetic basis of thermotolerance in *S. cerevisiae* with less attention devoted to cold tolerance, though numerous genes and pathways have been implicated [77–80,89–93]. Hybrids may offer a unique pathway for coping with temperatures above or below the optimal growing temperature of one parent [45,94,95], and may aid in the identification of genes important in temperature tolerance. For example, it has long been speculated that the allopolyploid hybrid yeast *S. pastorianus* (*S. cerevisiae* x *S. eubayanus*) tolerates the cold temperatures utilized in lager beer production due to the sub-genome of the cold adapted *S. eubayanus* [49–54,96–98]. Indeed, creation of *de novo* hybrids between *S. cerevisiae* and cold tolerant species *S. uvarum, S. eubayanus, S. arboricola*, and *S. mikatae* all show similar ability to ferment at 12°C [98]. A pair of recent studies show that mitochondrial inheritance in hybrids is also important in heat and cold tolerance, with the *S. cerevisiae* mitotype conferring heat tolerance and *S. uvarum* and *S. eubayanus* mitotypes conferring cold tolerance [95,99]. The hybrid ancestor used for our laboratory evolution experiments at both 15°C and 30°C has *S. cerevisiae* mitochondria, but exploring how this has influenced the evolution of these hybrids is worthy of further work.

Though our work here is complicated by utilizing multiple selection pressures (nutrient limitation and cold temperature), several patterns are suggestive of temperature specific adaptations in evolved hybrids. We see a slight single nucleotide variant bias towards *S. cerevisiae* mutations, we observe LOH events exclusively favoring the retention of the *S. uvarum* allele, and we demonstrate a fitness advantage of the *S. uvarum* allele compared to the *S. cerevisiae* allele at *PHO84*. The temperature sensitivity of the *PHO84* allele is a curious phenomenon for which we do not have a clear understanding. The phosphate metabolism pathway was recently implicated in temperature sensitivity in *S. cerevisiae* x *S. uvarum* hybrids: more specifically, *S. cerevisiae* alleles of 12 phosphate metabolism genes showed significantly higher expression at 37°C than *S. uvarum* alleles (no significant difference at 22°C or 33°C). *S. cerevisiae PHO84* was one of only a handful of genes whose expression was shown to be downregulated at 4°C and upregulated at 35°C [90]. These results are consistent with our observations from fitness assays that the *S. cerevisiae PHO84* allele is temperature sensitive, yielding high fitness at high temperatures. One potential connection is the need for inorganic phosphate for various processes involved in stress response, including heat shock and activation of the PKA pathway, of which *PHO84* is required [100–103]. This gene provides an interesting example of identifying a gene and pathway previously not appreciated for a role in temperature adaptation, and highlights using multiple environments to better understand parental species preferences and potentially environment specific incompatibilities.

More broadly, through the lens of *PHO84*, we establish LOH as an important molecular mechanism in hybrid adaptation, but we also demonstrate that this mutation type has fitness tradeoffs. The selection of a particular species allele may confer a fitness advantage in a given environment, but at a risk of extinction if the environment changes. Furthermore, such mutations rarely affect single genes, and instead operate on multigenic genomic segments, leading to a further pleiotropic benefit and/or risk even in environments unrelated to the initial selective regime. Relatively constant environments such as those found in the production of beer and wine may offer fewer such risks, where hybrids may find a particular niche that is less variable than their natural environment. Future efforts are warranted to explore how variable environments influence hybrid evolution and the extent of antagonistic pleiotropy in hybrid genomes. However, because LOH has been documented in a variety of different genera and taxa that experience a range of environments, it’s likely that our results have broad implications.

In conclusion, we illuminate pathways in which hybridization may allow adaptation to different temperature conditions. Mounting evidence suggests that anthropogenic climate change and habitat degradation are leading both to new niches that can be occupied by hybrids, as well as to new opportunities for hybridization due to changes in species distribution and breakdown of prezygotic reproductive isolation barriers [104–106]. Some researchers have speculated that this process is particularly likely in the arctic, where numerous hybrids have already been identified [107]. Our work supports the idea that portions of these hybrid genomes can be biased in parental representation by the environment in the initial generations following hybridization, and that this selection on species genetic variation may be beneficial or detrimental as conditions change.

## Materials and Methods

### Strains

Strains used to inoculate the laboratory evolution experiments and to gauge relative fitness of *PHO84* allele replacements in competition assays were previously utilized by Smukowski Heil et al. [60]. All strains are listed in **Table S4**.

### Evolution experiments

Continuous cultures were established using media and conditions previously described with several modifications to account for a temperature of 15°C [60,66]. Individual cultures were maintained in a 4°C room in a water bath such that the temperature the cultures experienced was 15°C, as monitored by a separate culture vessel containing a thermometer. The dilution rate was adjusted to approximately 0.08, equating to about 3 generations per day. Samples were taken twice a week and measured for optical density at 600 nm and cell count; microscopy was performed to check for contamination; and archival glycerol stocks were made. By 200 generations, 2/16 hybrid populations, 10/12 *S. cerevisiae* diploid populations, and 0/6 *S. uvarum* diploid populations had evolved a cell-cell sticking phenotype consistent with flocculation. The experiment was terminated at 200 generations and flocculent and non-flocculent populations were sampled from the final timepoint and submitted for whole genome sequencing (40 populations total, some cultures had only a flocculent or non-flocculent population while some cultures had both sub-populations). Populations from vessels that experienced flocculation were isolated as described in [84], and are denoted with “F”.

### Genome sequencing and analysis

DNA was extracted from each population using the Hoffman–Winston protocol (Hoffman and Winston 1987) and cleaned using the Clean & Concentrator kit (Zymo Research). Nextera libraries were prepared following the Nextera library kit protocol and sequenced using paired end 150 bp reads on the Illumina NextSeq 500 machine. The reference genomes used were *S. cerevisiae* v3 (Engel et al. 2014), *S. uvarum* (Scannell et al. 2011), and a hybrid reference genome created by concatenating the two genomes.

Variant calling was conducted on each population using two separate pipelines. For the first pipeline, we trimmed reads using trimmomatic/0.32 and aligned reads to their respective genomes (*S. cerevisiae., S. uvarum*, or a concatenated hybrid genome) using the mem algorithm from BWA/0.7.13, and manipulated the resulting files using Samtools/1.7. Duplicates were then removed using picard/2.6.0, and the indels were realigned using GATK/3.7. Variants were then called using Samtools (bcftools/1.5 with the–A and–B arguments), freebayes and lofreq/2.1.2. The variants were then filtered using bcftools/1.5 for quality scores above 10 and read depth above 20. For the second pipeline, reads were trimmed using Trimmomatic/0.32 and aligned using Bowtie2/2.2.3, then preprocessed in the same manner as the first pipeline. Variants were then called using lofreq/2.1.2 and freebayes/1.0.2–6-g3ce827d (using the –-pooled-discrete –-pooled-continuous –-report-genotype-likelihood-max –-allele-balance-priors-off –-min-alternate-fraction 0.05 arguments from bcbio (https://github.com/bcbio/bcbio-nextgen)). Variants were then filtered using bedtools/2.25.0 and the following arguments (Sup. table). In both variant calling pipelines, variants were filtered against their sequenced ancestors and annotated for gene identity, mutation type, and amino acid change consequence [108]. Final variant calls were manually confirmed through visual inspection in the Integrative Genomics Viewer [109] (1550 mutations checked in total).

For comparisons with clones evolved at 30°C which were analyzed using a different pipeline [60], we called variants on the previously published 30°C sequencing data using the same computational pipelines described here, and completely recapitulated the previous true positive variant calls.

### Data availability

Illumina reads generated in this study are deposited in the NCBI-SRA database under BioProject number PRJNA493117.

### Fitness assays

The pairwise competition experiments were performed in 20 ml chemostats as previously described [60,110]. The competition experiments performed at 15°C were modified as described above in the Evolution experiments. For all cold-evolved hybrid populations, one to two clones were isolated for use in competition experiments. Clones from P1–15° and P3–15° were PCR validated to have the chromosome XIII LOH event, but no other LOH, CNV, or single nucleotide variants were screened in these or any other clone tested.

## Supporting Information

**Table S1.** Mutations in cold-evolved *S. cerevisiae* diploid populations.

**Table S2.** Mutations in cold-evolved *S. uvarum* diploid populations.

**Table S3.** Comparison of single nucleotide variants called in 15°C and 30°C experimental evolution.

**Table S4.** Strain list.

**Figure S1.** Copy number plots of cold-evolved hybrids populations.

**Figure S2.** Amplification of *SUL1* in hybrids evolved at 15°C and 30°C.

**Figure S3.** Copy number plots of cold-evolved *S. cerevisiae* diploid populations.

**Figure S4.** Copy number plots of cold-evolved, flocculent *S. cerevisiae* diploid populations.

**Figure S5.** Loss of heterozygosity plots of cold-evolved *S. cerevisiae* diploid populations.

**Figure S6.** Loss of heterozygosity plots of cold-evolved, flocculent *S. cerevisiae* diploid populations.

**Figure S7.** Copy number plots of cold-evolved *S. uvarum* diploid populations.

**Figure S8.** Flocculation assay of several flocculent clones isolated from *S. cerevisiae* cold-evolved populations.

## Acknowledgements

We thank Angela Hickey for assistance with fitness assays. This work was supported by the National Science Foundation (grant number 1516330). M.J.D. is a Senior Fellow in the Genetic Networks program at the Canadian Institute for Advanced Research and a Rita Allen Foundation Scholar. M.J.D. is supported in part by a Faculty Scholars grant from the Howard Hughes Medical Institute.

## Author Contributions

C.S.H. and M.J.D. designed the experiment; C.S.H., C.R.L., and K.P. conducted experiments; C.S.H. and C.R.L. conducted data analysis; and C.S.H., C.R.L., and M.J.D. wrote the paper.

